# Oral mTOR Inhibition Limits And Reduces Actinic Keratosis And Cutaneous Squamous Cell Carcinoma In A UVB-Induced Mouse Model

**DOI:** 10.1101/2022.11.03.513568

**Authors:** MG Booty, B Komalo, A Hosny, SE Headland, MT Fernandez-Figueras, AM Nguyen, W Cousin, J Heinrich, L Nicolaisen, RM DeVay, B White, C Elabd

## Abstract

Actinic keratosis (AK) is a skin disease that is characterized by clinical and subclinical cutaneous lesions in sun-exposed areas. It is a considerable burden due to its high occurrence in middle-aged and older populations, as well as its propensity to progress to invasive cutaneous squamous cell carcinoma. The mammalian target of rapamycin (mTOR) pathway is critical in carcinogenesis and tumor development, and it has been shown to be over-activated during skin tumorigenesis, particularly upon ultraviolet (UV) radiation exposure, the key risk factor for AK. However, the ability of mTOR inhibitors to treat AK is not well documented. Herein, we evaluated the effect of oral mTOR inhibitors *in vitro* and *in vivo* and found that mTOR inhibitors lower keratinocyte cell proliferation *in vitro* and both clear and prevent AK and cutaneous squamous cell carcinoma (cSCC) in a UV-B induced SKH1 hairless mouse model of disease. mTOR inhibition reduced the number and size of skin lesions and the frequency of cSCC, resulting in a considerable reduction in disease severity. mTOR inhibition prevented lesion occurrence in areas of field cancerization, without affecting epidermal thickness, keratinocyte proliferation *in vivo*, or the presence of p53+ cells. Our findings indicate that, when appropriately dosed, oral mTOR inhibitors provide a safe home-based systemic treatment alternative with significant benefits to patients over current topical treatment options.

## INTRODUCTION

Actinic keratoses (AKs) are areas of slightly erythematous scaly skin that result from extended exposure to ultraviolet radiation. AKs are the most common precursors to cutaneous squamous cell carcinoma (cSCC), which is the second most common skin cancer globally ^1^. Though the majority of patients with cSCC have good prognoses, metastasis can occur, and due to the high incidence rate, it is estimated that cSCC has a death rate comparable to melanoma in the United States ^2^. The risk of a single AK progressing to cSCC is unclear but it is estimated to be as high as 16 % for immunocompetent patients ^3–7^.

The prevalence of AKs in the general population is high, particularly in individuals with a low skin phototype, and it increases with age. It therefore represents an important economical health care burden globally ^8,9^. Patients on immunosuppressive drugs have a higher prevalence of AKs and are at greater risk of progressing to cSCC ^4^. The estimated incidence of cSCC in immunosuppressed patients is 65-250 times higher than that of the general population ^10,11^. These observations are particularly relevant for solid organ transplant recipients who must remain on lifelong immunosuppressive drug regimens. However, not all immunosuppressive drugs are equivalent in terms of cancer risk. In kidney transplant recipients, maintenance immunosuppression with mammalian target of rapamycin (mTOR) inhibitors significantly reduces the risk of developing post-transplant malignancies ^12^. Specifically, mTOR inhibitors reduce the prevalence of skin cancers, including cSCC, and this effect is best observed when transplant recipients switch from calcineurin inhibitors to mTOR inhibitors early for maintenance immunosuppression ^13–17^.

The mechanisms underlying the reduced frequency of AKs and cSCC in transplant recipients treated with mTOR inhibitors remain unclear. Possibilities include the carcinogenic effects of calcineurin inhibitors and/or the impact of mTOR inhibition on keratinocyte growth and survival. Dysregulation of the PI3K/AKT/mTOR pathway is observed in multiple cancers and pathological skin conditions ^18^, and increased mTOR activity has been noted in AKs and cSCC ^19,20^. Enhanced mTOR activity is broadly associated with the growth and survival of transformed cells ^21^, thus, mTOR inhibition may represent a therapeutic intervention capable of impacting both AKs and cSCC. In a transgenic mouse model of cSCC, topical administration of the mTOR inhibitor BEZ235 significantly reduced tumor size ^22^, and mTOR inhibition is reported to limit the growth and survival of human cSCC and keratinocyte cell lines ^23–25^. These studies suggest mTOR inhibition may impact AKs and cSCC by limiting keratinocyte growth and survival in these lesions.

Histological analysis can be used to assess lesion severity and likelihood of progression into cSCC. AK is typically initiated by a transformation of the basal layer of the epidermis that results in the proliferation of abnormal keratinocytes. Microscopically, the basal layers show a loss of polarization with nuclear pleomorphism and hyperchromasia, as well as an increase in the mitotic activity and superficial hyperkeratosis. In addition, there is often a lymphocytic infiltrate of variable intensity ^26^. Early AK with a transformed basal compartment can lead to one of three fates. It can remain stable, regress, or progress. Progression can lead directly to the development of an invasive cSCC or result in a complete turnover of the normal keratinocytes within the epidermis before evolving into an invasive cSCC ^26^. For practical purposes, the progressive replacement of the epidermis has been arbitrarily divided into three stages, classified as AK-I, AK-II, and AK-III. AK-I corresponds to lesions with atypia limited to the lower third of the epidermis, AK-II is used when the atypical features involve the two lower thirds of the epidermis, while AK-III is used when the upper third is involved. This later stage is also described as in situ cSCC. When the epidermal/dermal junction is disrupted, then it can be considered that an invasive cSCC has developed ^27,28^. Cumulative sun damage can lead to the development of mutation-rich areas in the skin through a process known as field cancerization. These areas are susceptible to the development of AK and subsequent progression to cSCC. Histologically, the cancer field shows atypia and disorder at the basal layer, similar to early AK. The absence of hyperkeratosis, acanthosis or inflammation explains the clinical invisibility.

We hypothesize that mTOR inhibition can reduce and prevent AKs and limit their progression to cSCC. Systemic administration of mTOR inhibitors has the added advantage of targeting all clinical and subclinical keratoses, and it differs from current standards of care that rely on lesion-directed procedures or topical field therapies. By simultaneously treating all AKs, the incidence of cSCC could be dramatically reduced. Systemic mTOR inhibition has been tested in multiple clinical trials seeking to improve immune function in the elderly and can be well tolerated when dosed appropriately ^29–31^. For these reasons, we examined the impact of oral mTOR inhibition in a mouse model of UV-induced AK and cSCC.

In this study, we demonstrated that systemic mTOR inhibition effectively reduces keratinocyte cell growth *in vitro* and rapidly clears and prevents AK and cSCC in UV-B induced SKH1 hairless mice. Moreover, mTOR inhibition decreased the number and size of skin lesions and reduced the frequency of cSCC, leading to an overall significant decrease in disease severity. mTOR inhibition led to the prevention of lesion formation in areas of field cancerization; however it did not affect epidermal thickness, keratinocyte proliferation *in vivo* or the presence of p53+ cells in the epidermis upon UV-B exposure.

## RESULTS

To assess the impact of mTOR inhibition on keratinocyte growth and survival *in vitro*, we subjected the human keratinocyte cell line, HaCaT, to varying concentrations of mTOR inhibitors. The HaCaT cell line is derived from adult trunk skin that was spontaneously immortalized and has UV-indicative p53 mutations found in both cSCC and normal human skin ^32,33^. The three mTOR inhibitors selected for testing were vistusertib (AZD2014), omipalisib (GSK2126458), and CC-115, which are all dual mTORC1 and mTORC2 inhibitors ^34–37^. In addition to targeting mTOR, omipalisib is a potent PI3K inhibitor, and CC-115 inhibits DNA-PK. Proliferation was assessed by measuring confluence of cells plated at a low density over time in a live cell imaging chamber. All three mTOR inhibitors demonstrated dose-dependent inhibition of keratinocyte proliferation in concentrations ranging from 39 nM to 10 µM (**Figure 1a**). Importantly, we did not observe cell loss in these cultures so changes in confluency are likely attributable to anti-proliferative mechanisms and not cell death. In order to test the effect of mTOR inhibitors on cellular survival, apoptotic cell death was measured on fully confluent (non-proliferative) HaCaT cell cultures. Keratinocyte apoptosis was quantified by counting cells with active caspase-3 and -7 over 48 hours of drug exposure. Only omipalisib increased the number of apoptotic cells over DMSO when dosed at concentrations ranging from 39nM to 10 µM (**Figure 1b**). As a proof of concept, these experiments indicate that vistusertib, omipalisib, and CC-115 can inhibit keratinocyte proliferation and may be suitable to treat or slow the development of actinic keratosis (AK).

**Figure 1:**
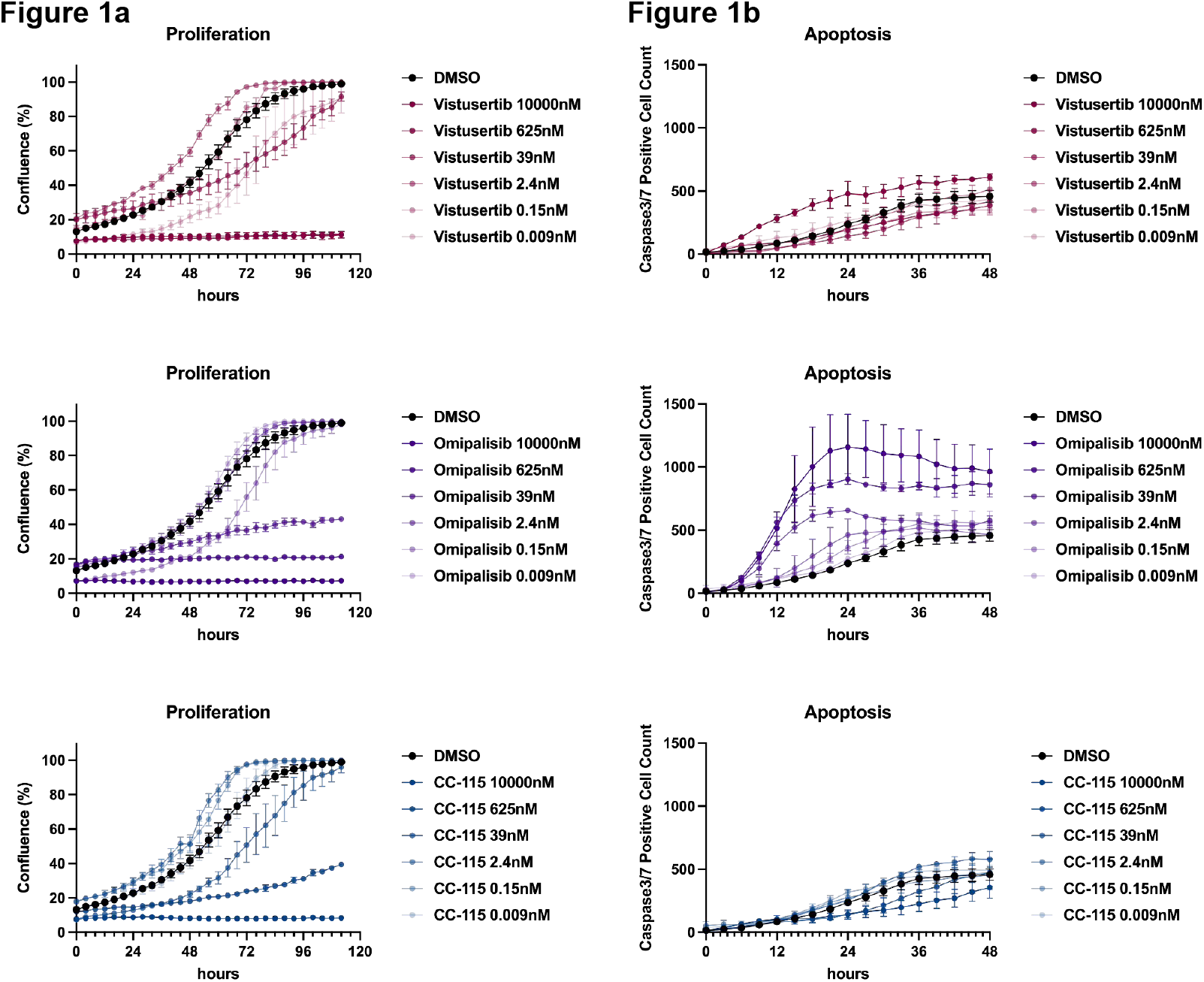
mTOR Inhibition slows keratinocyte cell growth in vitro. HaCat cells were plated in 96 well plates at low density (2,500 cells/well) for cell proliferation assays or at high density (10,000 cells/well) for apoptosis assays. Cells were exposed to various concentrations of mTOR inhibitors as indicated. **Figure 1a**. Proliferation, presented as the percentage of confluence, was determined every 4 hours using the Incucyte® live-cell analysis systems. **Figure 1b**. Apoptosis, presented as the number of caspase 3/7 positive cells, was quantified every 3 hours using the Incucyte® live-cell analysis systems. Data represent the mean +/- SEM of 2 independent experiments.

UV-B irradiated SKH1 hairless mice are the gold standard preclinical model to study the development of AK and cSCC. Because this mouse system recapitulates many characteristics of human disease ^38^, it constitutes a predictive model to identify novel therapeutic options for both early and advanced stages of AK and cSCC ^38,39^. The efficacy of many approved treatments for AK and cSCC (e.g. 5-Fluorouracil, photodynamic therapy, ingenol mebutate and diclofenac) has previously been demonstrated using this model ^40–43^.

**Figure 2a** details the *in vivo* experimental design used to examine mTOR inhibition. SKH1 hairless mice were exposed to UV-B radiation 3 times per week for a total of 16 weeks. The incrementally increased radiation levels and exposure schedule are described in **Supplemental Table 1**. This protocol was successfully used in previous studies ^38^ and generates a broad spectrum of lesion severity ranging from early AK to cSCC **(Figure 2b**). Following UV-B exposure, animals were rested for one week before randomization into the control group (n = 16 mice) and the test article groups (n = 10 mice per group). Animals were randomized based on the presence of lesions to ensure an even distribution across the groups at the pre-treatment baselines. At randomization, 62.8% of the animals presented one or more skin lesions. Lesion counts and body weight were balanced between the groups (**Supplemental table 2**).

**Figure 2:**
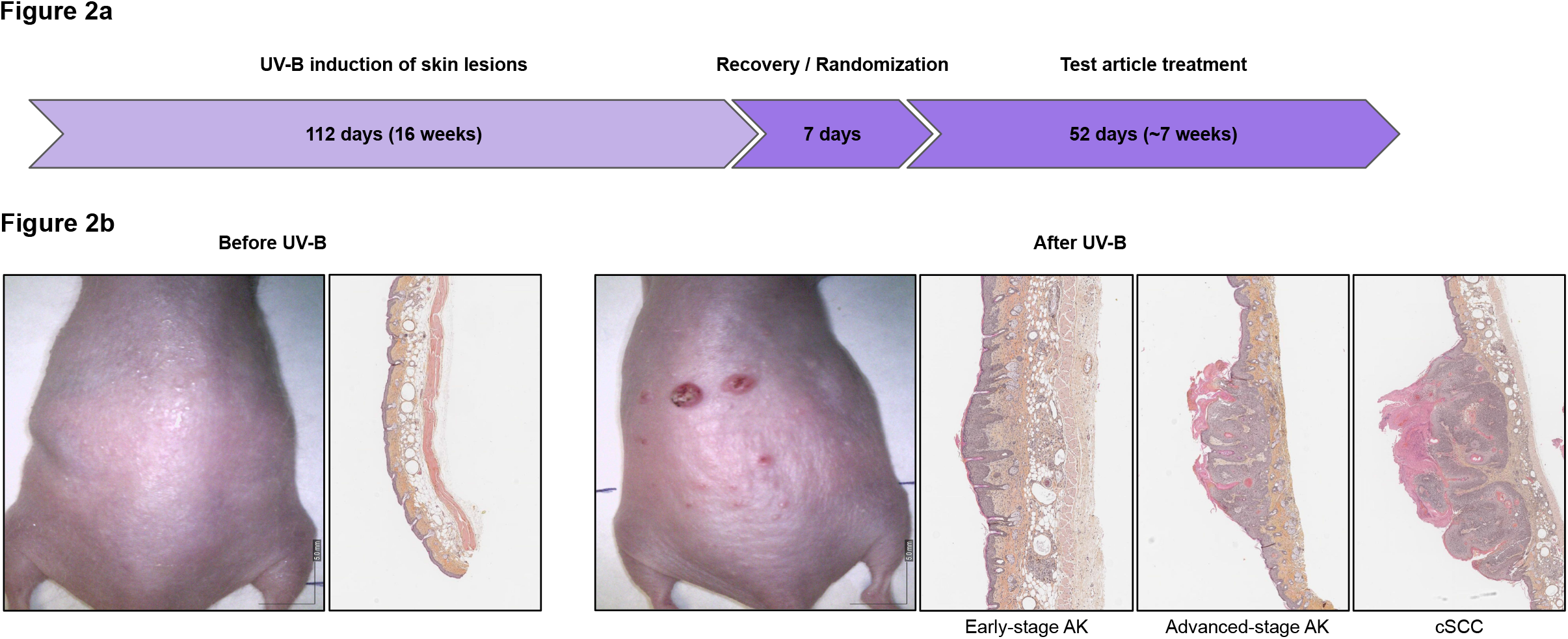
SKH1-Elite hairless mouse model for AK and cSCC. SKH1-Elite hairless mice were exposed to UV-B radiation for 16 weeks (112 days) in order to develop AK and cSCC. Animals recovered from UV-B radiation exposure for 7 days before vehicle or test article treatments were administered orally for a duration of 7 weeks (52 days). **Figure 2a**. Experimental design schematic representation. **Figure 2b**. Representative dermatoscope images and skin histology (hematoxylin eosin and saffron staining) of non-UV-B exposed (left) and UV-B exposed (right) mice. Early-stage AK, advanced-stage AK and cSCC are present in UV-B-exposed mice.

After the 16-week UV-B induction of lesions, mice were subjected to test article treatments via oral gavage. To test the impact of mTOR inhibition in the treatment of AK and cSSC, three mTOR inhibitors were dosed orally once daily for a duration of 52 days. Prior to study start, separate pharmacokinetic studies were performed for all compounds following single-dose oral administration to better inform the study design. All three mTOR inhibitors showed robust systemic exposure following oral administration. The pharmacokinetic parameters are listed in **Supplemental table 3**. Of the three compounds, vistusertib exhibited the highest *C*_*max*_ and *AUC* and omipalisib had the lowest values. Despite having lower overall serum concentrations, omipalisib has the lowest reported IC_50_ for mTOR (∼10 fold lower than vistusertib and ∼70 fold lower than CC-115), thus its activity is likely to remain high at relatively low doses ^34–37^.

In a previously conducted experiment using the same animal model, we observed that vistusertib was well tolerated at a dose of 15 mg/kg and appeared to limit the development of keratoses (data not shown). In this experiment, vistusertib was therefore administered at a dose of 15 mg/kg. In addition to pharmacokinetic analyses, a tolerability study was performed to determine suitable long-term dosing regimens for omipalisib and CC-115. In healthy mice dosed daily for 14 days, omipalisib was tolerated at doses up to 6 mg/kg and CC-115 at doses up to 10 mg/kg without inducing clinical adverse events (mortality, abnormalities or signs of pain) or weight loss (**Supplemental table 4**). Based on pharmacokinetics, tolerability, and published literature, doses for omipalisib were chosen to be 0.1 and 1 mg/kg, and doses for CC-115 were chosen to be 1 and 5 mg/kg.

Representative images of the different groups after 52 days of treatment are presented in **Figure 3a. Figure 3b** displays weekly images of a representative animal throughout its treatment course. A timelapse of 9 animals from the vehicle control group, the vistusertib (15 mg/kg) group and the omipalisib (1 mg/kg) group is also presented in **Supplemental Figure 1**. Visual inspections revealed that both vistusertib and omiplasib treated animals had reduced lesions relative to the vehicle control group at the end of treatment. CC-115 treatment groups also had fewer lesions, though the effects were the least pronounced compared to the other mTOR inhibitors **(Figure 3a)**. As shown in **Figure 3b**, mice generally displayed a few keratoses of a moderate size for the first 10 days of treatment. As time progressed, the number and size of keratoses increased in vehicle-treated mice despite the absence of UV-B exposure, indicating that the cellular damage that accumulated during UV-B exposure continued to drive AK formation. mTOR inhibition both resolved pre-existing keratoses and prevented formation of new ones over time **(Figure 3b, c, and d)**. At the end of the treatment phase, only 20% of the mice in the vistusertib group and 10% of the mice in the omipalisib high-dose group displayed 5 or more keratoses compared to ∼44% of the mice in the vehicle group. In addition, none of the lesions present in these mTOR inhibitor groups reached the size of the largest lesions observed in the vehicle group **(Figure 3 and Supplemental Figure 1)**.

**Figure 3:**
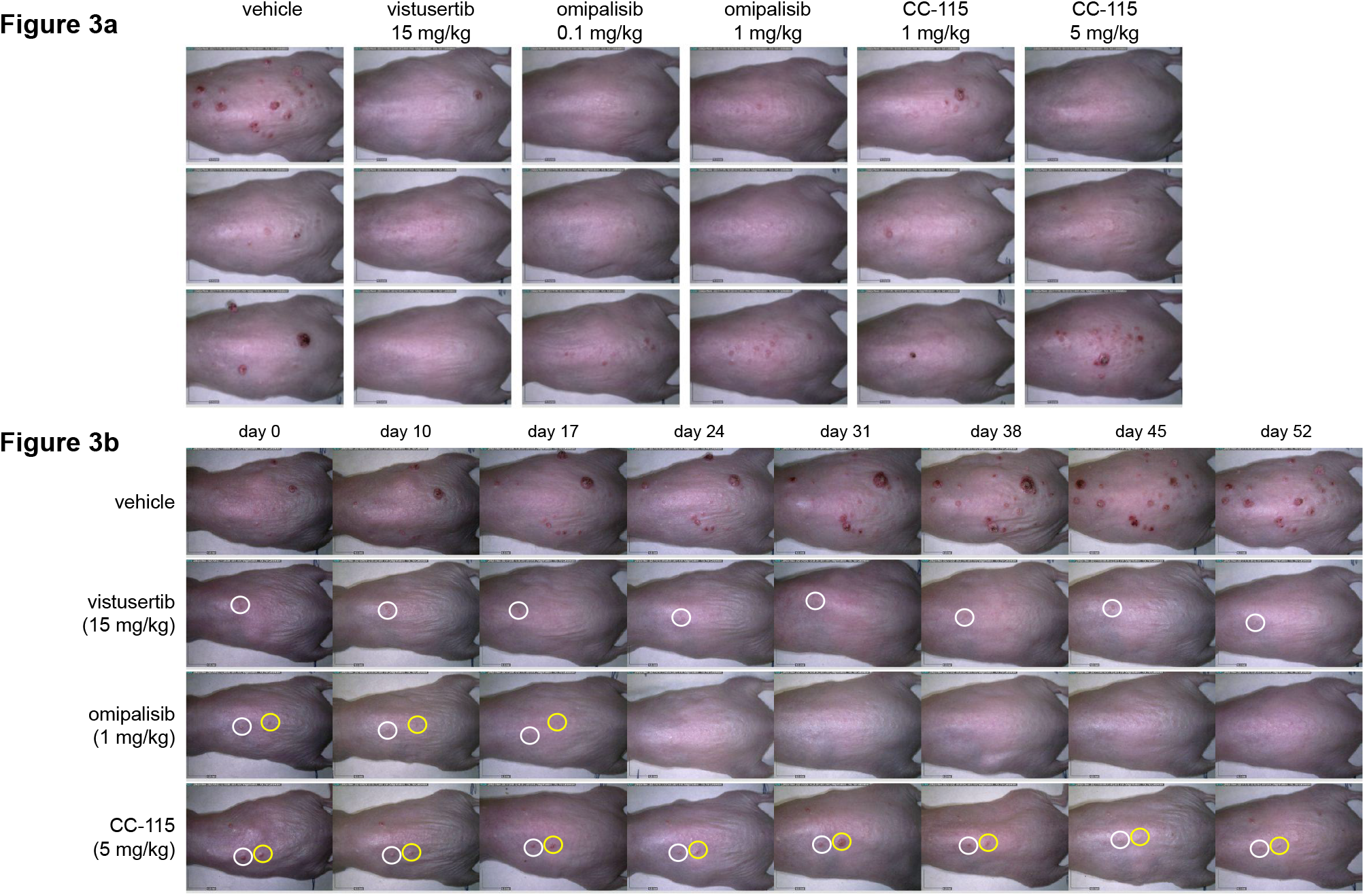
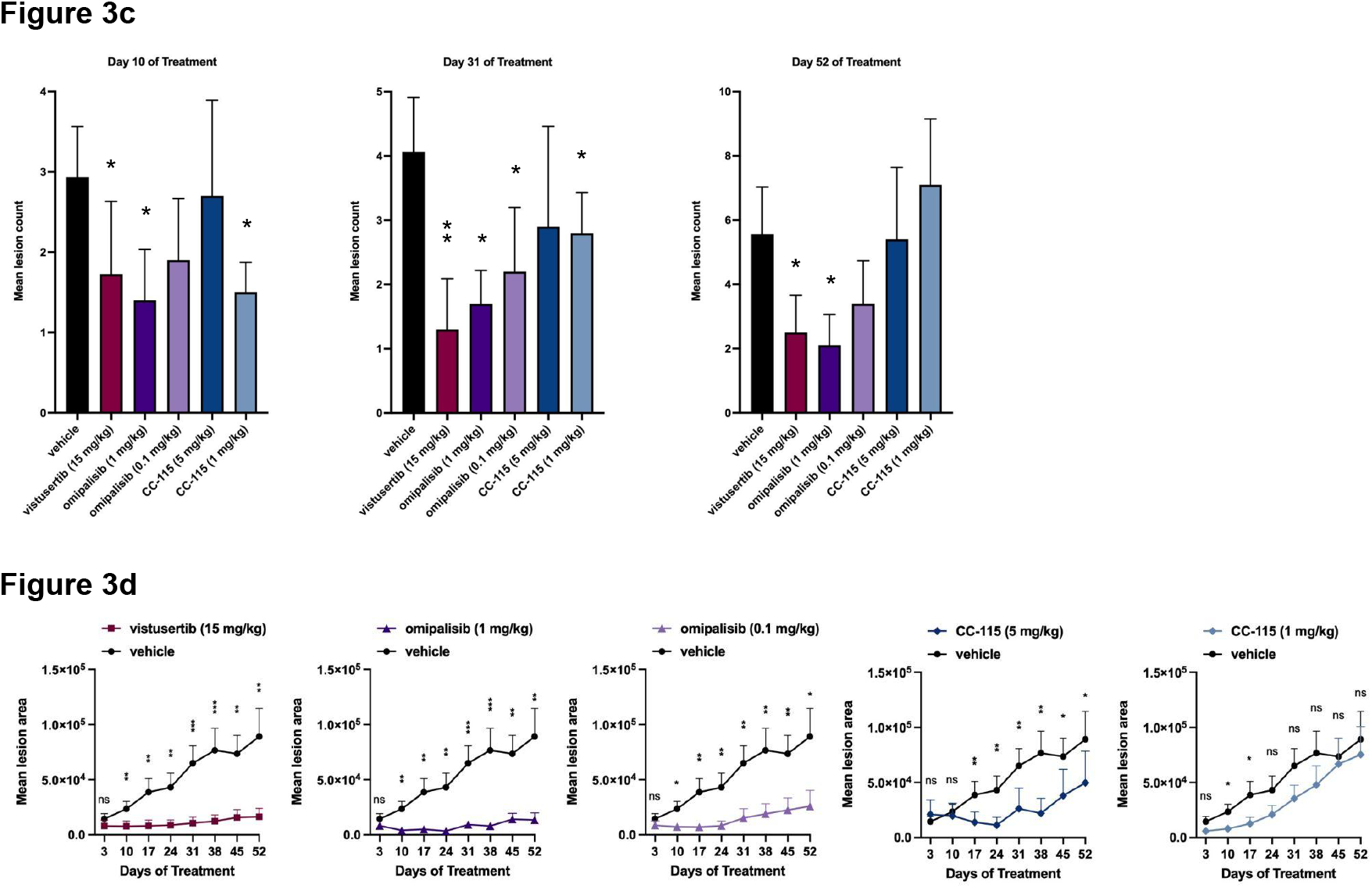
mTOR inhibition reduces lesion number and size in the UV-B induced SKH1-Elite hairless mouse model. Dermatoscope images were acquired at randomization and weekly during the 52 days of test article treatment. **Figure 3a**. Dermatoscope images of 3 representative mice for each group after 52 days of vehicle or test article treatment. **Figure 3b**. Time course dermatoscope images of one representative mouse from vehicle, vistusertib (15 mg/kg), omipalisib (1 mg/kg) and CC-115 (5 mg/kg) groups. White and yellow circles track lesions over time and show lesion clearance upon mTOR inhibition for lesions present prior to treatment initiation. Lesions increased in number and size over time in the vehicle control group; they tend to scab and fall out towards day 45 or day 52 leaving a clear lesion on the skin. **Figure 3c**. Bar graphs represent the average number of skin lesions per mouse for each group after 10, 31 and 52 days of vehicle or test article treatment. Lesions were counted on dermatoscope images in a blinded manner. Data represents mean +/- SEM (n = 16 mice for the vehicle control, n = 9 mice for CC-115 (1 mg/kg) at day 31 and 52, and n = 10 mice for all other test article groups). Differences in lesion counts between test article groups and the vehicle control group were determined using a two-tailed Mann-Whitney U test, *\* P value <0*.*05, ** P value < 0*.*01*. **Figure 3d**. Lesion size measured on dermatoscope images after 3, 10, 17, 24, 31, 38, 45 and 52 days of vehicle or test article treatment. The average total lesion area is presented for each group compared to vehicle control. Data represent mean total lesion area +/- SEM (n = 16 mice for the vehicle control, n = 9 mice for CC-115 (1 mg/kg) at day 31 and 52, and n = 10 mice for all other test article groups). Differences in lesion size between test article groups and the control group were determined using a two-tailed Mann-Whitney U test, *\* P value <0*.*05, ** P value < 0*.*01* and *\*\*\* P value < 0*.*001*.

Lesion quantification for each group was performed on dermatoscope images collected weekly for each mouse during the treatment regimen, and all lesions with a diameter equal to or greater than 1 mm were included in the lesion counts. As shown in **Figure 3c**, the vistusertib (15 mg/kg) treated mice showed significantly reduced lesion numbers by day 10, and this reduction persisted throughout the treatment course. Both high- (1 mg/kg) and low- (0.1 mg/kg) dose omipalisib groups had fewer lesions by day 31; however, only high-dose omipalisib maintained a significant reduction in the number of lesions by the end of the experiment. CC-115 had the lowest impact on lesion counts, with the low dose (1 mg/kg) showing a reduction in lesion counts early on that failed to persist to the end of treatment.

Vistusertib and omipalisib treated groups displayed a significant reduction in lesion size as determined by their mean total lesion area compared to the vehicle group. These differences were significant as early as 10 days after treatment initiation (**Figure 3d**). Calculated total lesion area remained significantly lower than the vehicle control for these treatment groups for the entire 52 days of treatment. Despite minimal differences in lesion counts, high-dose CC-115 (5 mg/kg) showed a significant decrease in lesion size compared to the control group by day 17 and this difference persisted for the duration of the treatment course. For low-dose CC-115 (1 mg/kg), a minor reduction in lesion area was noted on days 10 and 17, but this was not maintained throughout the treatment course. For all successful treatment groups, therapeutic efficacy was associated with preventing the formation of new lesions, restraining the growth of existing lesions, and fully resolving some lesions.

After 52 days of test article treatments, animals were euthanized and all lesions from each mouse were collected and processed for histopathological assessment. Skin lesions were fixed, paraffin embedded and stained with haematoxylin, eosin and saffron (HES) and sent for blinded evaluation by a histopathologist. The decrease in lesion counts observed upon mTOR inhibition (**Figure 3c**) is accompanied by a change in the proportion of the different types of lesions. mTOR inhibition strongly shifted the proportion of lesions classified as cSCC to less severe, lower grades. 36.1% (30/83) of the lesions in the vehicle group were classified as cSCC, whereas the proportion of cSCC was reduced to 21.4% (6/28), 25% (5/20) and 18.8% (6/32) in mice treated with vistusertib, high-dose omipalisib (1 mg/kg) and low-dose omipalisib (0.1 mg/kg), respectively (**Figure 4a**). Though CC-115 did not significantly reduce the number of lesions after 52 days of treatment, it reduced the proportion of lesions classified as cSCC. cSCC represented 25% of the lesions of mice treated with high-dose CC-115 (5 mg/kg) and 28% of the lesions of mice treated with low-dose CC-115 (1 mg/kg). The decrease in the proportion of cSCC in the vistusertib and the omipalisib high-dose (1 mg/Kg) groups was accompanied by an increase in the proportion of AK-I (vistusertib = 17.9%; omipalisib high-dose = 15%; vehicle = 8.4%) and AK-II (vistusertib = 28.6%; omipalisib high-dose = 35%; vehicle = 22.9%) lesions compared to the vehicle group. The decrease in the proportion of lesions classified as cSCC in the omipalisib low-dose (0.1 mg/kg) and the CC-115 high-dose (5 mg/kg) groups was accompanied by an increase in the proportion of AK-III lesions (omipalisib low-dose = 46.9%; CC-115 high-dose = 44.4%; vehicle = 32.53%) compared to the vehicle group (**Figure 4a**).

**Figure 4:**
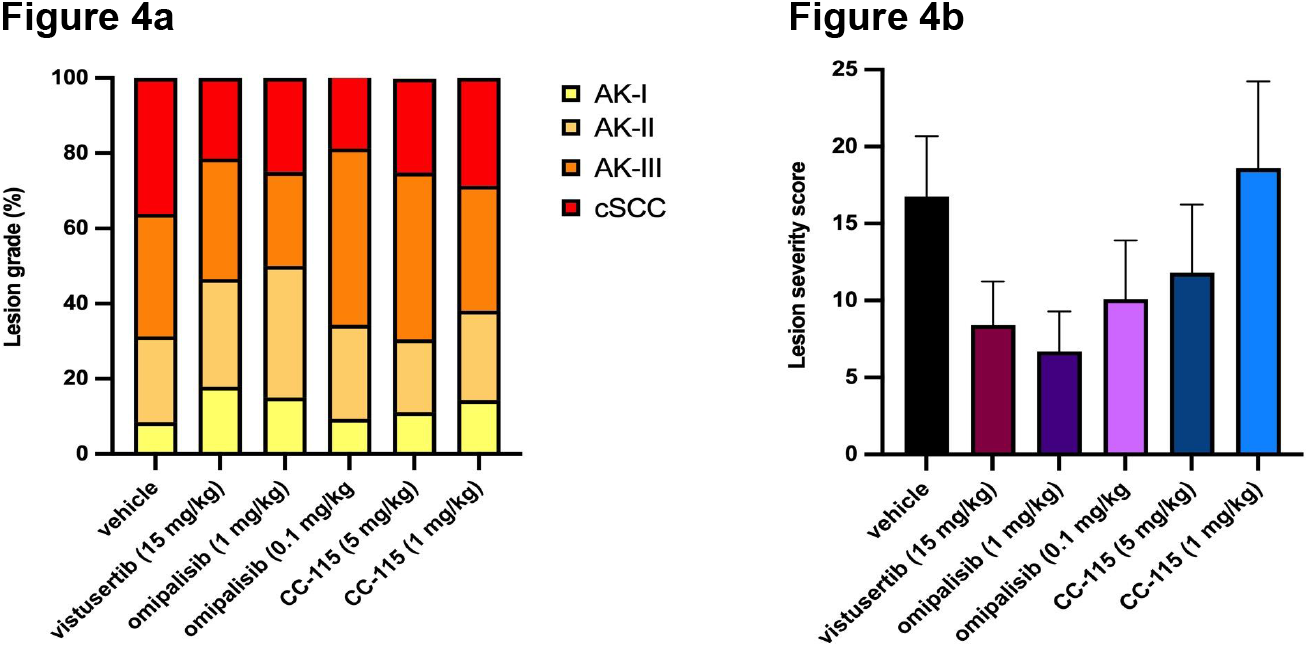
mTOR inhibition reduced lesion severity and decreased the occurrence of cSCC. Histopathological grading of lesions collected after 52 days of vehicle or test article treatment was performed. **Figure 4a**. Proportion of AK-I, AK-II, AK-III and cSCC for each group. **Figure 4b**. Lesion severity score for each group. Data represents mean +/- SEM (n = 16 mice for the vehicle control, n = 9 mice for CC-115 (1 mg/kg), and n = 10 mice for all other test article groups).

To get a complete picture of treatment efficacy, we combined lesion numbers and lesion histopathological grading to create a severity score. Each lesion was first assigned a score based on their histopathological grading (AK-I = 1; AK-II = 2; AK-III = 3, cSCC = 4). A lesion severity score was then calculated for each mouse by cumulating their lesions scores (severity score = number of AK-I *1 + number of AK-II*2 + number of AK-II *3 + number of cSCC*4). The severity score for each group was finally calculated by averaging individual mouse scores for each group (**Figure 4b**). Importantly, all three mTOR inhibitors showed remarkable reduction in the group lesion severity score compared to the vehicle group.

In the UV-B irradiated SKH1 hairless mouse model, there are relatively few lesions present at the end of the UV-B irradiation phase (average of 2 lesions per mouse) compared to 52 days after UV-B is stopped (average of ∼5 lesions per mouse) (**Figure 3c and Supplemental Table 2**). Thus, the majority of lesions present in the vehicle group at day 52 developed as time progressed during the treatment phase. The efficacy of mTOR inhibitors to reduce lesion number therefore primarily reflects their ability to prevent new lesions from forming or progressing in the cancer field. To test this hypothesis we performed epidermal thickness measurements and immunohistochemistry on paraffin embedded sections of skin taken from all animals at day 52. The analysis was performed using unbiased and automated quantification as described in **Supplemental Figure 2**.

For epidermal thickness measurements, a deep learning model was trained and validated for the automated segmentation of epidermal tissue on HES-stained slides as described in the methods (**Figure 5a and Supplemental Figure 3**). As expected, UV-B exposure induced a 2 fold increase in the epidermal thickness over non-UV-B treated animals (**Figure 5b**). While no significant differences in epidermal thickness of the cancer field was observed between the vehicle group and mTOR inhibitor treated groups (**Figure 5b**), a positive correlation between epidermal thickness and lesion counts was observed (**Supplemental Figure 4a**).

**Figure 5:**
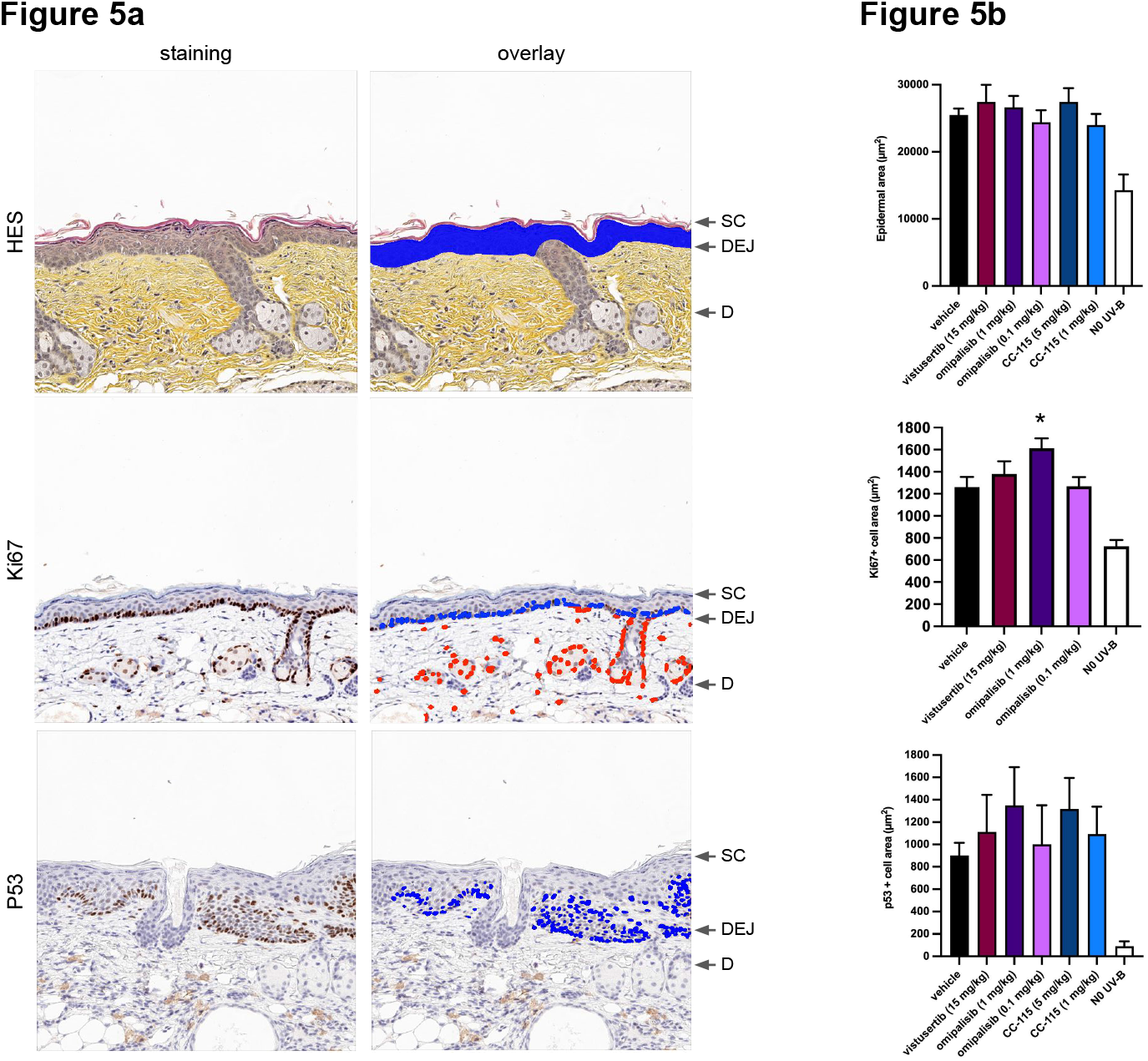
mTOR inhibition does not impact epidermal thickness, Ki67, or P53 in the cancer field. Skin tissue samples were collected from areas of field cancerization 52 days after vehicle or test article treatment. **Figure 5a**. Skin tissue sections were stained with hematoxylin eosin and saffron (HES, top), for Ki67 (middle) or for P53 (bottom). Stained sections were scanned with a high resolution scanner and tiled. Automated segmentation of the epidermis is shown in blue overlay in the top right panel, automated detection of Ki67+ cells in the epidermis (blue) and in the dermis (red) is shown in the middle right panel and automated detection of P53+ cells in the epidermis is shown in blue in the bottom right panel. **Figure 5b**. Quantification of epidermal thickness (top), keratinocyte proliferation (Ki67+ cell area, middle), and P53+ cell area (bottom) are shown for the indicated groups. Data represents mean +/- SEM (n = 16 mice for the vehicle control, n = 9 mice for CC-115 (1 mg/kg), and n = 10 mice for all other test article groups). Two-tailed Mann-Whitney U test, *\* P value <0*.*05*. D: dermis; DEJ: dermal-epidermal junction; SC: stratum corneum.

Immunohistochemical analysis was next performed to test for the presence of Ki67+ and p53+ cells in the epidermis. Similar to our epidermal thickness analysis, we developed automated pipelines for detecting and segmenting cells that were positive for these molecular targets (**Supplemental Figure 2**). It is well established that an increase in the proliferation of keratinocytes (Ki67+) in the cancer field, particularly in regions that are distal from the dermal-epidermal junction (DEJ), correlates with an increased risk of developing keratosis ^44^. **Figure 5a** shows representative Ki67 immunostaining of a cancer field section taken from a vehicle control animal. Overlays indicate automated detection of Ki67+ cells in the epidermis and the dermis. As expected, UV-B exposure increased the proportion of proliferating (Ki67+) keratinocytes in the epidermis of UV-B exposed animals relative to non-UV-B exposed animals (**Figure 5b**). In this dataset, there was a positive correlation between keratinocyte proliferation and lesion counts (**Supplemental Figure 4b**). However, no difference in the proportion of Ki67+ cells were observed upon mTOR inhibition with the exception of a minor increase in the omipalisib (1 mg/kg) treatment group.

It is well established that UV-B irradiation induces mouse skin cancers containing p53 mutations at a high frequency, and that these mutations precede the appearance of skin tumors ^45^. **Figure 5a** shows representative p53 immunostaining and the automated detection of p53+ cells. A significant increase in p53+ cells was observed in the vehicle group compared to mice not exposed to UV-B (**Figure 5b**). A strong positive correlation was observed between p53 and Ki67 (**Supplemental Figure 4c**). However, no difference in the proportion of p53+ cells was observed between the vehicle group and any of the mTOR inhibitor groups. Overall, these data demonstrate that mTOR inhibition is efficacious in treating and preventing AKs and cSCC in the cancer field independently of cell growth inhibition or the clearance of cells accumulating p53 mutations.

## DISCUSSION

Mammalian target of rapamycin (mTOR) is an evolutionarily conserved protein kinase that regulates growth (mass and size), metabolism and cell division in all eukaryotic cells by incorporating nutrient and mitogen signals ^46,47^. mTOR complex 1 (mTORC1) mainly regulates cell growth and metabolism, while mTOR complex 2 (mTORC2) mainly controls cell proliferation and survival ^48^. Our findings are in line with the role of mTOR controlling cellular proliferation in human keratinocytes with all three mTOR inhibitors tested demonstrating a dose dependent inhibition of cellular proliferation. Interestingly, omipalisib was the only mTOR inhibitor to induce apoptosis in human keratinocytes. While all three mTOR inhibitors tested are dual mTORC1 and mTORC2 inhibitors, omipalisib is additionally a potent phosphatidylinositol-3-kinase (PI3K) inhibitor which could explain its ability to induce apoptosis ^49,50^.

mTOR plays an important role in tumorigenesis and tumor development with multiple studies describing that tumors over-activate the PI3K/AKT/mTOR signaling pathway ^21,51^. mTOR pathway over-activation has also been observed during skin tumorigenesis and in particular upon UV-irradiation. In this context, unbalanced AKT/mTOR signaling may eventually lead to hyper-proliferation and contribute to malignant transformation ^52^. A previous report demonstrated the efficacy of the mTOR inhibitor BEZ235 in reducing tumor size in the K14-FYN-Y528F transgenic mouse model when applied topically ^22^. These mice develop spontaneous cSCC via constitutive activation of the Src-family kinase Fyn in basal keratinocytes. In the present study we report similar efficacy observations. We showed for the first time the efficacy of three mTOR inhibitors in treating and preventing AK and cSCC upon systemic administration in a physiological animal model, *i*.*e*. long-term low-level UVB-exposure that recapitulates many characteristics of human disease ^38^. Hence, our data open the possibility of using self-administered systemic treatments as a safe option with lower patient-burden than existing treatments.

Ideally, actinic keratosis treatment should remove as many clinical and subclinical lesions as possible, achieve a long-lasting clinical remission, produce a pleasing cosmetic outcome, and stop the development of invasive squamous cell carcinoma ^53^. Current standards of care for AK patients rely on lesion-directed procedures or topical field therapies. Topical field therapies require multiple weeks of application per treatment area and oftentimes patients have to sequentially rotate treatment of multiple affected areas leading to extended treatment periods. Patients frequently struggle to apply topical therapies properly and cope with undesirable local skin reactions, resulting in less than optimal treatment adherence. A systemic approach could alleviate many shortcomings associated with topical field therapies by simultaneously treating clinical and subclinical lesions from all the affected fields. Unfortunately, to date, no oral treatment is approved for AK or cSCC.

Data presented herein support the use of mTOR inhibitors administered orally for the treatment of AK with the potential to dramatically reduce the incidence of cSCC. In cancer patients, systemic mTOR inhibition has been associated with adverse side effects and increased risk of hyperglycemia, hypertriglyceridemia, hypercholesterolemia, thrombocytopenia, anemia, nausea, and stomatitis ^54,55^. However, in multiple clinical trials seeking to improve immune function in the elderly, systemic mTOR inhibition has been tested and was well tolerated when dosed appropriately ^29–31^. Safety in elderly populations is key since advanced age is one of the major risk factors for AK and cSCC ^56^, and it can be achieved by lowering the dose or adapting the dosing regimen, maintaining efficacy with minimal side effects. The mTOR pathway regulation involves multiple feedback mechanisms that may be differentially activated depending on the degree of mTOR inhibition. In a recent study, Antoch MP *et al*. ^57^ showed higher efficacy of the lower dose of a highly bioavailable mTOR inhibitor in the prevention of prostate cancer in a transgenic mouse model ^57^.

In the UV-B irradiated SKH1 hairless mouse model, the majority of lesions present in the vehicle group at the end of the in-life assessment (day 52) developed post UV-B exposure, as time progressed during the treatment phase of the study. The efficacy of mTOR inhibitors to reduce lesion number therefore primarily reflects their ability to prevent or treat subclinical lesions. We also studied the effect of mTOR inhibitors on well established readouts in the cancer field. Specifically, measured epidermal thickness and quantified the presence of Ki67 or p53 positive cells in the epidermis. Interestingly, mTOR inhibitors did not reduce the increased epidermal thickness typically associated with UV-B exposure. These results are in accordance with Ki67 quantifications that showed no difference in the number of proliferating (Ki67+) keratinocytes between the vehicle control group and the mTOR treated groups. If anything, a small increase in the number of Ki67+ cells was observed upon omipalisib exposure at 1 mg/kg. Mutations in the TP53 gene typically alter p53 protein degradation, therefore p53 presence and staining intensity is often associated with the presence of p53 mutations leading to cancer formation. As expected UVB increased the number of p53 positive cells in the cancer field but mTOR inhibitors had no effect.

Overall, these data demonstrate that mTOR inhibition is efficacious in treating and preventing AK and cSCC independently of cell growth inhibition or the clearance of cells accumulating p53 mutations. The mechanism of action of mTOR inhibition in the context of UV-B-induced AK and cSCC remains to be elucidated and further exploration is needed.

## MATERIALS AND METHODS

### Cell Culture

HaCaT cells were cultured in growth medium composed of CnT-PR (Zen-Bio) and penicillin-streptomycin (500 IU ml^-1^, 0.1 mg ml^-1^; Gibco) on plates coated with recombinant human collagen I (Coating Matrix kit, Gibco) according to manufacturer’s instructions. Cells were maintained in 5% CO2, 95% humidity and 37 degree celsius. Twenty four hours before an experiment, cells were washed with Hanks Balanced Salt Solution (HBSS, Gibco) and passaged using TrypLE-select (Gibco). Cells were plated into 96 well flat bottom plates in growth medium supplemented with 10 µM of ROCK inhibitor (Stem Cell Technologies). Cells were plated at low density (2,500 cells/well) for cell proliferation assays or at high density (10,000 cells/well) for apoptosis assays. The ROCK inhibitor was only present for 16-24 hours after plating. On the day of the experiment, cells were exposed to various concentrations of mTOR inhibitors or DMSO vehicle control (Sigma-Aldrich) diluted in culture medium and imaged over time using the Incucyte® Live-Cell Analysis Systems (for confluence). For the apoptosis assay, Caspase 3/7 indicator dye (Sartorius, Germany) and propidium iodide (BD Biosciences) were added to the culture medium before mTOR inhibitor treatment. Images were analyzed using the supplied manufacturer software and confluency (% of the surface area covered by cells) and the number of caspase 3/7 positive cells were quantified.

### Animals

Six-week-old female SKH1-Elite hairless mice were purchased from the Charles River Laboratories. Animals were acclimated for six days upon receipt at the animal facility. Animals were housed in shoe-box cages with filter tops and in a room dedicated to mice. General procedures for animal housing and husbandry were conducted according to the testing facility standard operating procedures and met all regulations concerning use of animals in research including the U.S. Department of Agriculture regulations (9 CFR Ch.1) implementing the Animal Welfare Act (7 USC 2131 et seq.) and the recommendations of the National Research Council’s “Guide for Care and Use of Laboratory Animals” (National Academy Press, 1996). Mice were provided ad libitum water and food (LabDiet® 5001 Rodent Diet, Purina Mills, Inc., St. Louis, MO) throughout the acclimation and in-life phases and were kept under a twelve hours of light and twelve hours of dark cycle with controlled temperature ranging from 20-26 degree celsius and humidity between 30 and 70%. All procedures were conducted in accordance with the Institutional Animal Care and Use Committee of the testing facility. Sample size for each experiment was determined based on previously published differences between treated and untreated UV-B exposed mice in order to have sufficient statistical power, using the minimum necessary number of mice.

Actinic keratoses and cSCC were induced upon 16 weeks of UV-B exposure. UV-B radiation was performed 3 times per week following radiation levels and the schedule described in **Supplemental Table 1**. This protocol was successfully used in previous studies ^38^ and generates a broad spectrum of lesion severity ranging from early AK to cSCC (**Figure 2b**). Following UV-B exposure, animals were rested for one week before randomization into the control group (n = 16 mice) and the test article groups (n = 10 mice per group). Animals were randomized based on the presence of lesions to ensure an even distribution across the groups at the pre-treatment baselines.

mTOR inhibitors were diluted in DMSO (Sigma-Aldrich) and stored at -80 degree celsius as concentrated aliquots. Prior to dosing, aliquots were thawed and formulated in 0.5% methylcellulose and 10% DMSO. mTOR inhibitors were administered orally, daily for up to 52 days.

### Reagents

mTOR inhibitors vistusertib (AZD2014) (CAS No. 1009298-59-2), CC-115 (CAS No. 1228013-15-7) and omipalisib (CAS No. 1086062-66-9) were purchased from MedChemExpress. Anti-Ki67 rabbit polyclonal antibody (ab15580) was purchased from Abcam and was used at a 1:800 dilution. Anti-p53 rabbit polyclonal antibody (clone CM5) was purchased from Leica Biosystem (NCL-L-p53-CM5p) and was used at a 1:1000 dilution.

### Histology and immunohistochemistry

Skin samples were collected on day 52 of test article treatment after mice were euthanized. All lesions plus one cancer field skin sample were collected from each mice included in the study. Skin samples were fixed in 10% neutral buffered formalin and embedded in paraffin. 5 µm thick sections were collected and stained with haematoxylin, eosin and saffron (HES) for histopathological assessment. Immunohistochemistry analysis was performed on 5 µm sections stained with anti-Ki67 rabbit polyclonal antibody or anti-p53 rabbit polyclonal antibody (clone CM5) and counterstained with haematoxylin. Following primary antibody incubation, a secondary HRP-conjugated antibody was used and signals was revealed following DAB addition.

### Computational histopathological analysis

The computational image analysis workflow for calculating epidermal thickness and immunohistochemistry analysis is illustrated in Supplemental Figure 2. All histopathology sections were digitally acquired.

### Data preprocessing and quality control

The dermal-epidermal junction (DEJ) was manually delineated on all sections. Each section was then fragmented into 512 × 512 μm square tiles aligned to the DEJ, generating from 12 to 25 tiles per section. A subset of these tiles was manually labeled for the absence or presence of epidermal tissue and were subsequently used to train a convolutional neural network model ^58^ to classify tiles accordingly. The trained convolutional neural network model was used to perform automated quality control where tiles without epidermal tissue were excluded.

### Epidermal thickness

Epidermal thickness was quantified on haematoxylin, eosin and saffron (HES) stained sections. Manual segmentation of the epidermal tissue was performed in a random subset of 515 tiles equally stratified across donors and groups. Among these 515 manually segmented tiles, 412 were used to train a fully convolutional neural network to automatically segment epidermal tissue. Testing of this trained algorithm was performed in the hold-out set (n=103) and it scored a volumetric dice of 0.92, a high geometric overlap with 1 being full overlap. See **Supplemental Figure 3** for examples of manual and automated epidermis segmentations. The validated algorithm was then used to perform epidermis segmentation on all tiles. Epidermal thickness was calculated as the epidermal area based on the automated segmentations, divided by DEJ length.

### Immunohistochemistry analysis

Ki67 or p53 positive signals were quantified on immuno-histochemistry (IHC) stained sections using the open-source software CellProfiler™ (www.cellprofiler.org). Segmentation of Ki67 and p53 positive pixels was performed using manually predefined imaging features and the area of Ki67 or p53 positive pixels for each tile was quantified. Data aggregation per animal was performed by averaging positive pixel area from all the tiles extracted from the animals sections.

## ACKNOWLEDGEMENTS

We thank Ben Kamens and Spring Discovery, Inc. for funding this work.

