## Supplemental Material for "Oral mTOR Inhibition Limits And Reduces Actinic Keratosis And Cutaneous Squamous Cell Carcinoma In A UVB-Induced Mouse Model"

### TITLE

Short Title:
Oral mTOR inhibition treats actinic keratosis

### AUTHORS

Booty MG, Komalo B, Hosny A, Headland SE, Fernandez-Figueras MT, Nguyen AM, Cousin W, Heinrich J, Nicolaisen L, DeVay RM, White B, Elabd C*.

***Correspondence**
Christian Elabd, PhD
1121 Industrial Road, Suite 500, San Carlos, CA 94070
+1 510 224 4484


### SUPPLEMENTARY MATERIAL

**Supplemental Table 1: UV-B radiation exposure**
Mice were exposed to UV-B radiation 3 times per week for a total of 16 weeks in order to develop AK and cSCC. The minimal erythema dose (MED) was identified as 500 J/m2 for SKH1-Elite hairless mice. UV-B radiation exposure was performed as indicated.

**Supplemental Table 2: Lesion counts and body weights at randomization**
After 16 weeks of UV-B exposure, mice were randomized based on lesion counts. No statistical differences in lesion counts and body weights were observed at randomization using a two-tailed Mann–Whitney U test, n=16 mice for the control group and n=10 mice per treatment group.

**Supplemental Table 3: mTOR inhibitors pharmacokinetics**
Single doses of vistusertib (15 mg/kg), omipalisib (3 mg/kg) and CC-115 (1 mg/kg) were administered orally using the same formulation as used in the study (10% DMSO and 0.5% methylcellulose). Blood was collected at 0.5, 1, 2, 4, 8 and 24 hours following administration using $K_2EDTA$ as anticoagulant. Vistusertib, omipalisib and CC-115 were measured in plasma samples using validated mass spectrometry methods. Pharmacokinetic parameters are

summarized. AUC: area under the curve; Tmax: time to reach maximum concentration; Cmax: maximum concentration.

**Supplemental Table 4: mTOR inhibitors maximum tolerated dose**
Vistusertib, omipalisib and CC-115 were administered orally daily at various concentrations over a 14-day period using the same formulation used in the study (10% DMSO and 0.5% methylcellulose). The maximum tolerated dose summarized in the table corresponds to the maximum daily dose tested that was well tolerated in mice as determined by the absence of clinical observations (mortality, abnormalities, signs of pain) and body weight loss inferior or equal to 5% from baseline.

**Supplemental Figure 1: Dermatosocpe images timelapse**
Dermatoscope images timelapse of mice before and weekly after vehicle control, vistusertib (15 mg/kg) or omipalisib (1 mg/kg) treatments. Nine representative mice are shown for each group.

**Supplemental Figure 2: Computational histopathology workflow**
Diagram outlining the computational workflow. First, the dermis-epidermis junction (DEJ) is manually identified on the sections. 512 pixel by 512 pixel tiles are then extracted along the DEJ with a scale of 1μm equals to 1 pixel. Tiles without epidermis are filtered out through an automated quality control step. Image quantification was performed on the tiles in order to calculate the epidermis area, measure Ki67 positive cell area, and measure p53 positive cell area.

**Supplemental Figure 3: Manual versus automated epidermis segmentation**
Skin cross sections were stained with hematoxylin eosin and saffron and the epidermis tissue was either manually segmented (top row) or automatically segmented (bottom row) using a machine learning model trained on manually annotated data.

**Supplemental Figure 4: Correlations between lesion counts and cancer field features**
Correlations between lesion count and epidermal area (**Supplemental Figure 4a**), lesion count and Ki67+ area (**Supplemental Figure 4b**), as well as Ki67+ area and p53+ area (**Supplemental Figure 4c**) are shown. Correlation was measured using the Pearson correlation coefficient.
