## Supplemental Tables and Figures for "Oral mTOR Inhibition Limits And Reduces Actinic Keratosis And Cutaneous Squamous Cell Carcinoma In A UVB-Induced Mouse Model"

### Slide 1
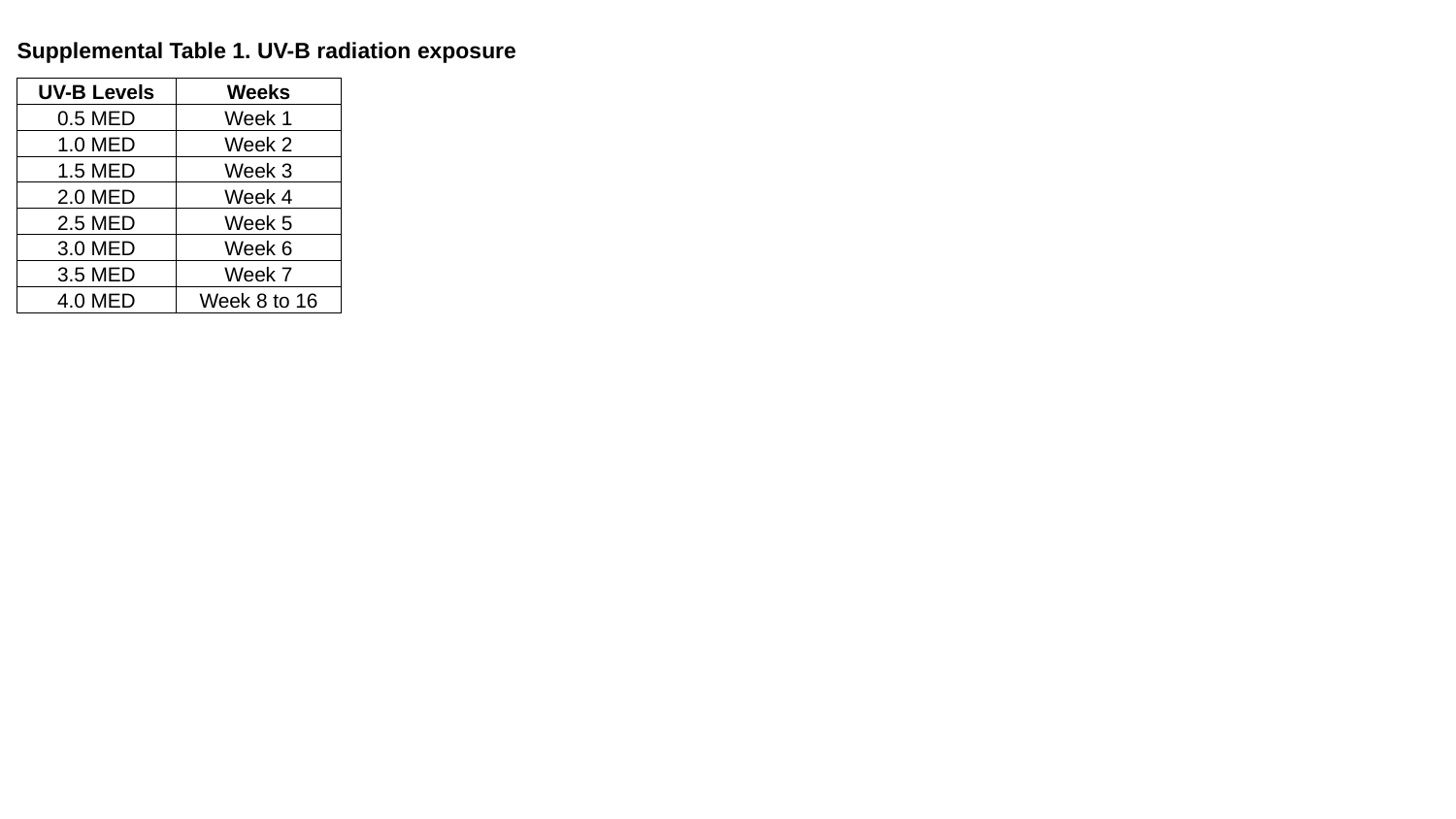

Supplemental Table 1. UV-B radiation exposure
| UV-B Levels | Weeks |
| --- | --- |
| 0.5 MED | Week 1 |
| 1.0 MED | Week 2 |
| 1.5 MED | Week 3 |
| 2.0 MED | Week 4 |
| 2.5 MED | Week 5 |
| 3.0 MED | Week 6 |
| 3.5 MED | Week 7 |
| 4.0 MED | Week 8 to 16 |

### Slide 2
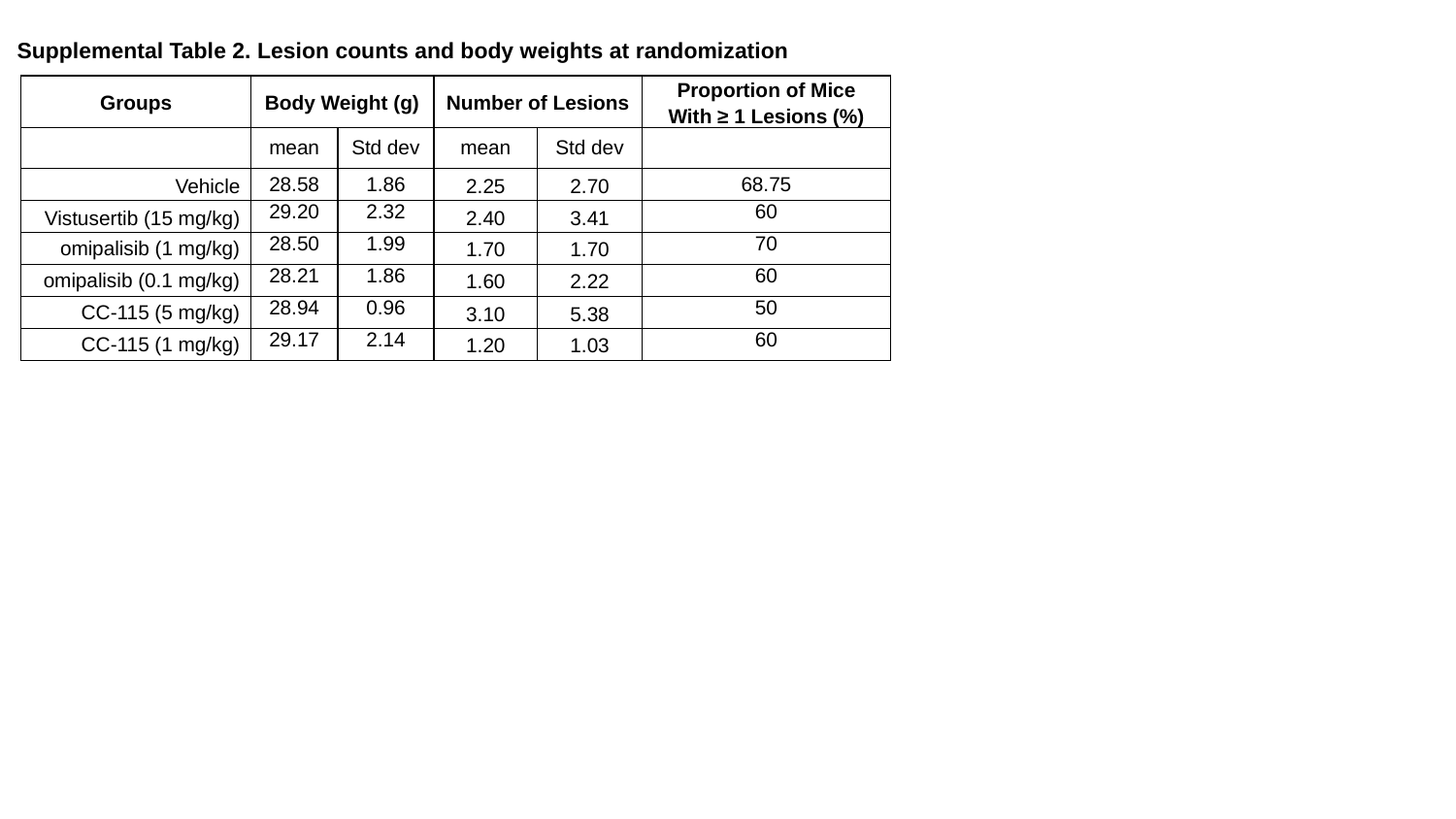

Supplemental Table 2. Lesion counts and body weights at randomization
| Groups | Body Weight (g) | | Number of Lesions | | Proportion of Mice With ≥ 1 Lesions (%) |
| --- | --- | --- | --- | --- | --- |
| | mean | Std dev | mean | Std dev | |
| Vehicle | 28.58 | 1.86 | 2.25 | 2.70 | 68.75 |
| Vistusertib (15 mg/kg) | 29.20 | 2.32 | 2.40 | 3.41 | 60 |
| omipalisib (1 mg/kg) | 28.50 | 1.99 | 1.70 | 1.70 | 70 |
| omipalisib (0.1 mg/kg) | 28.21 | 1.86 | 1.60 | 2.22 | 60 |
| CC-115 (5 mg/kg) | 28.94 | 0.96 | 3.10 | 5.38 | 50 |
| CC-115 (1 mg/kg) | 29.17 | 2.14 | 1.20 | 1.03 | 60 |

### Slide 3
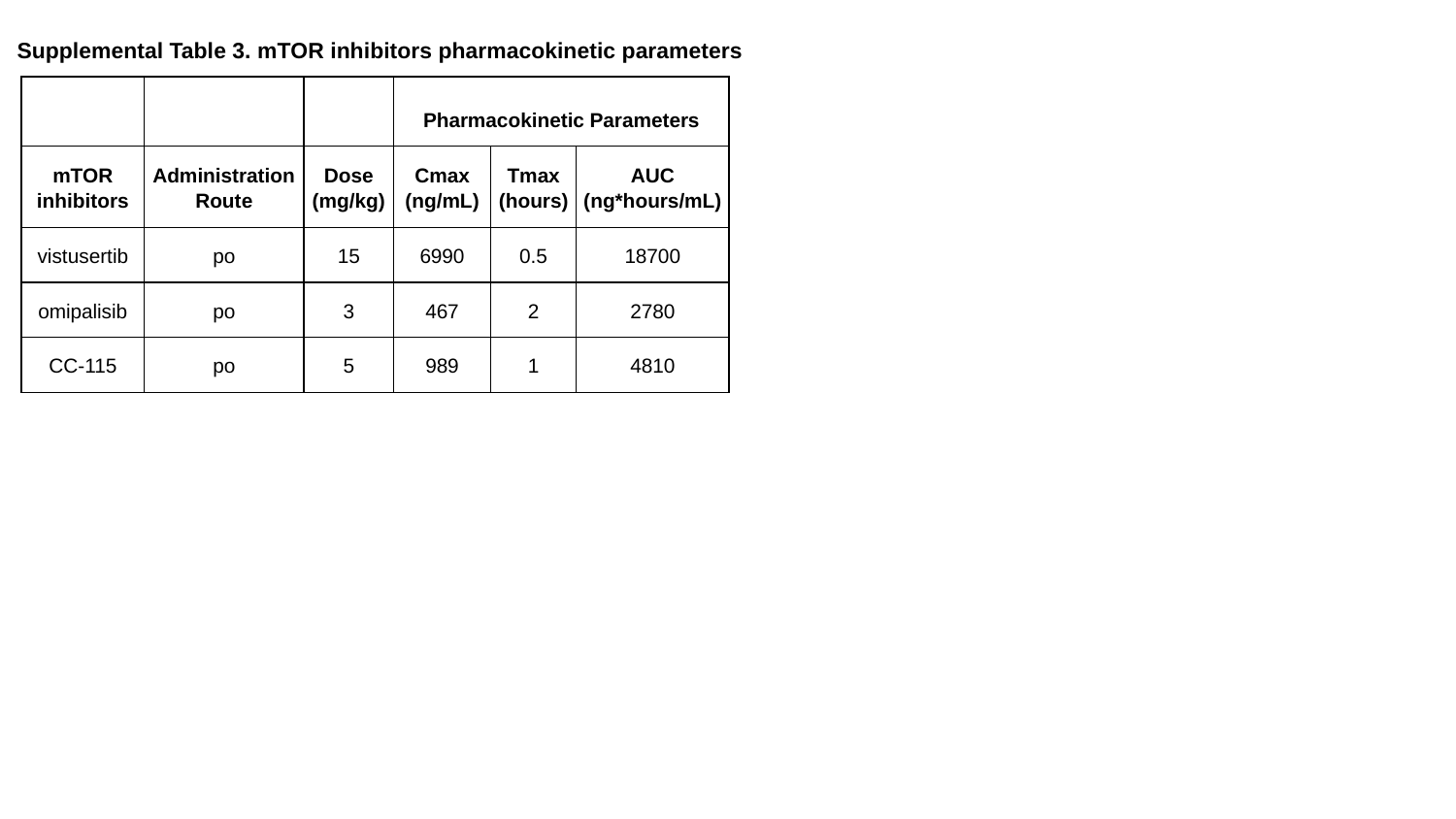

Supplemental Table 3. mTOR inhibitors pharmacokinetic parameters
| | | | Pharmacokinetic Parameters | | |
| --- | --- | --- | --- | --- | --- |
| mTOR inhibitors | Administration Route | Dose (mg/kg) | Cmax (ng/mL) | Tmax (hours) | AUC (ng\*hours/mL) |
| vistusertib | po | 15 | 6990 | 0.5 | 18700 |
| omipalisib | po | 3 | 467 | 2 | 2780 |
| CC-115 | po | 5 | 989 | 1 | 4810 |

### Slide 4
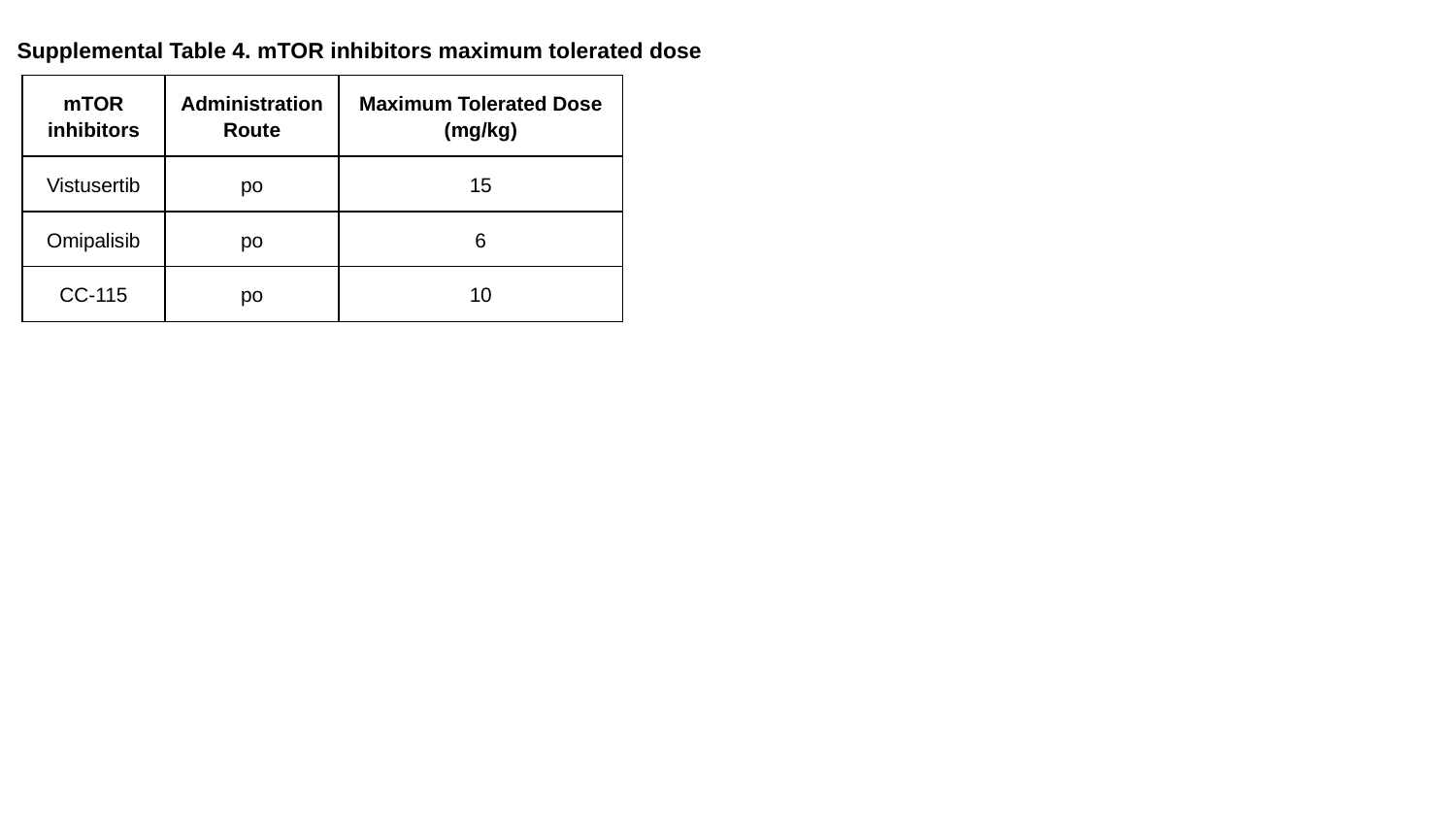

Supplemental Table 4. mTOR inhibitors maximum tolerated dose
| mTOR inhibitors | Administration Route | Maximum Tolerated Dose (mg/kg) |
| --- | --- | --- |
| Vistusertib | po | 15 |
| Omipalisib | po | 6 |
| CC-115 | po | 10 |

### Slide 5
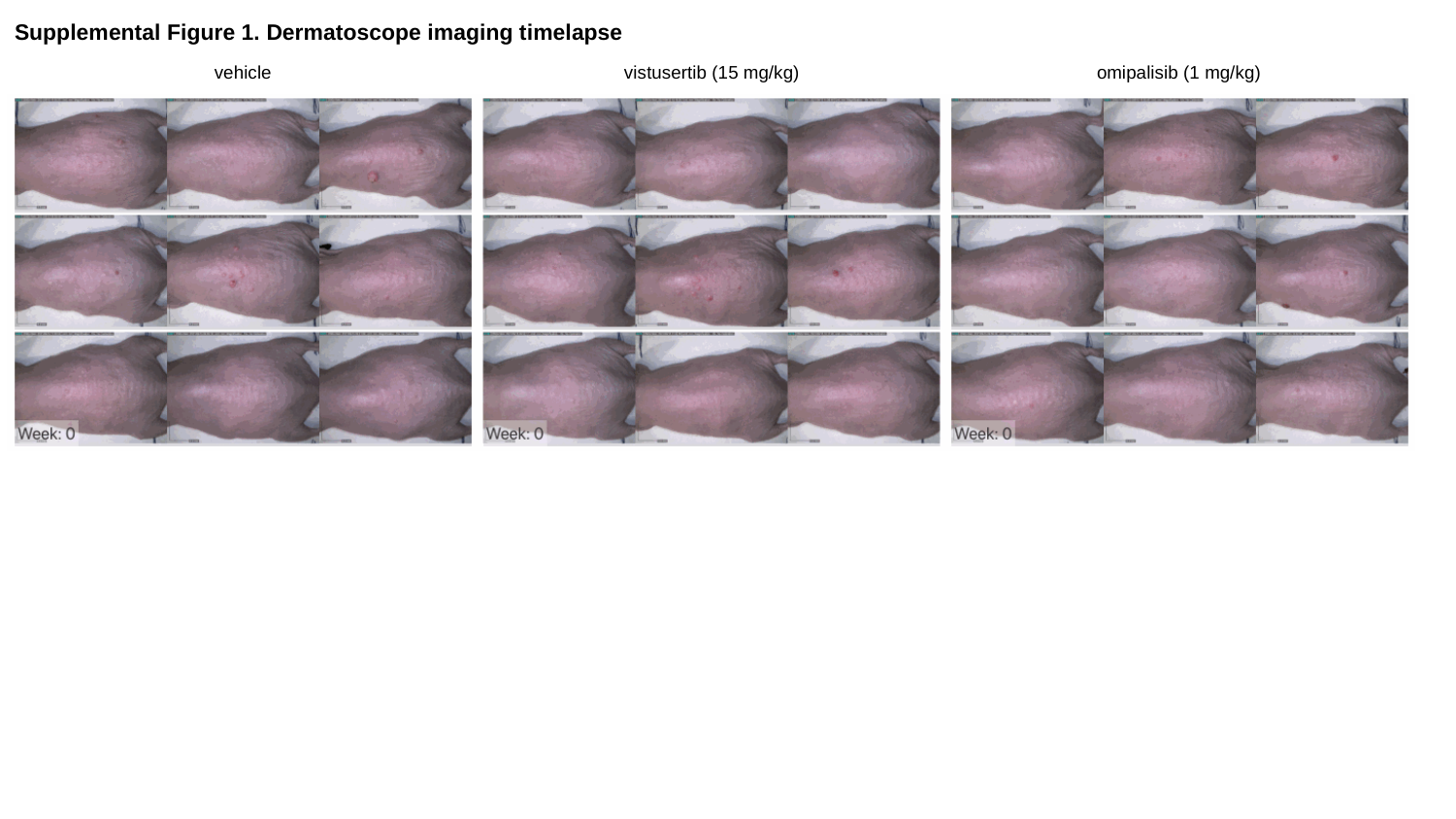

Supplemental Figure 1. Dermatoscope imaging timelapse
vistusertib (15 mg/kg)
omipalisib (1 mg/kg)
vehicle

### Slide 6
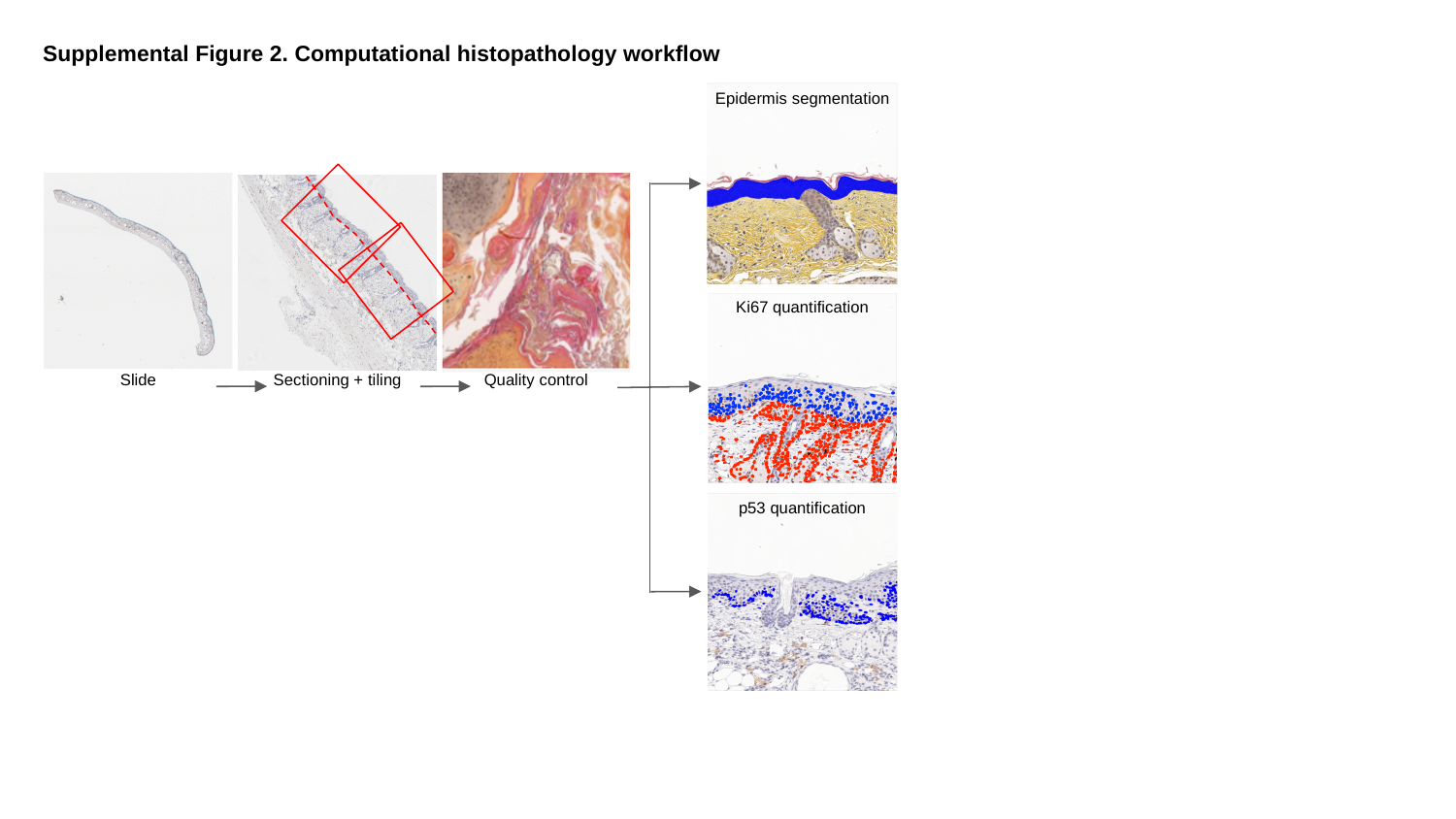

Supplemental Figure 2. Computational histopathology workflow
Epidermis segmentation
Ki67 quantification
Sectioning + tiling
Slide
Quality control
p53 quantification

### Slide 7
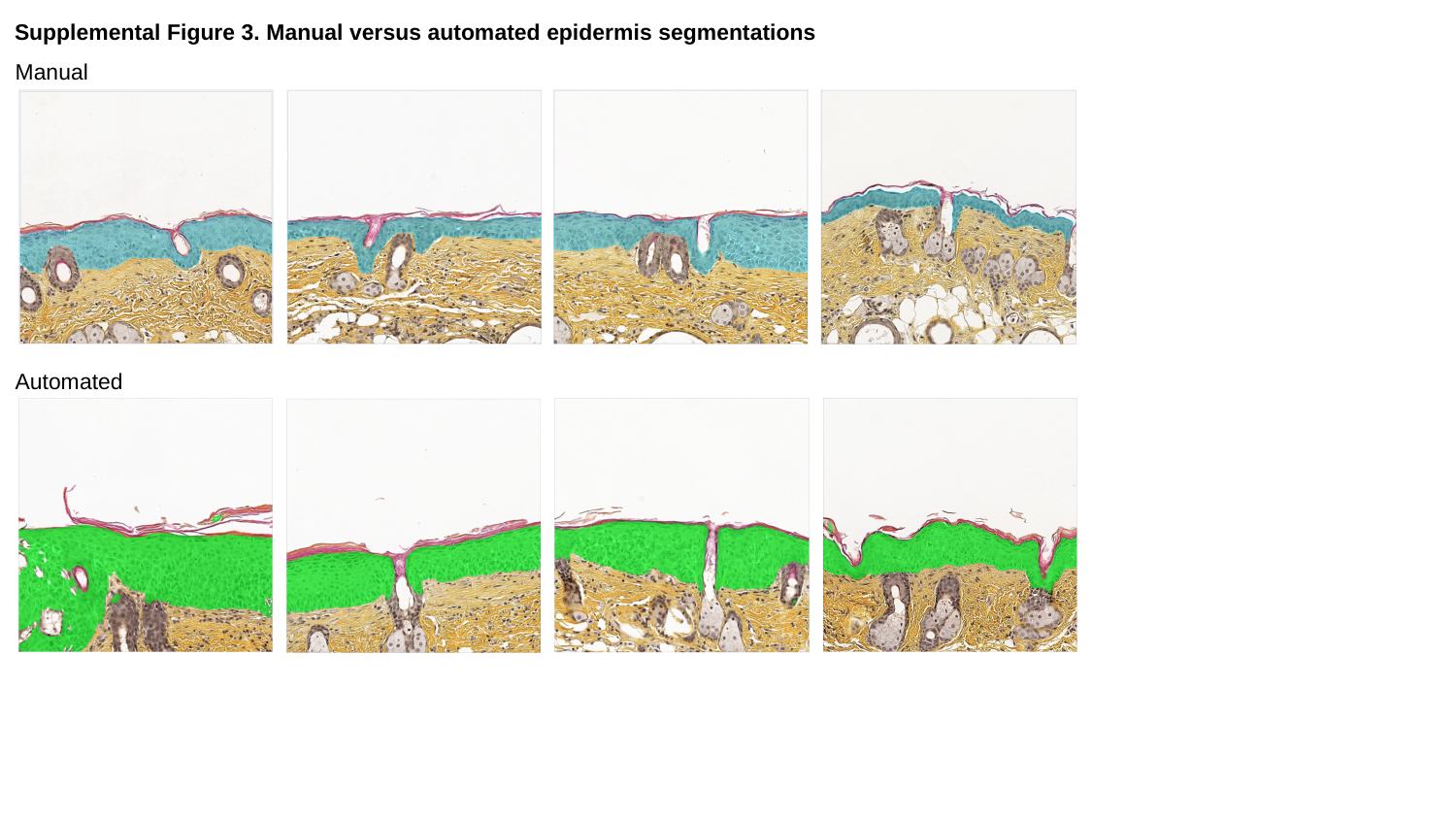

Supplemental Figure 3. Manual versus automated epidermis segmentations
Manual
Automated

### Slide 8
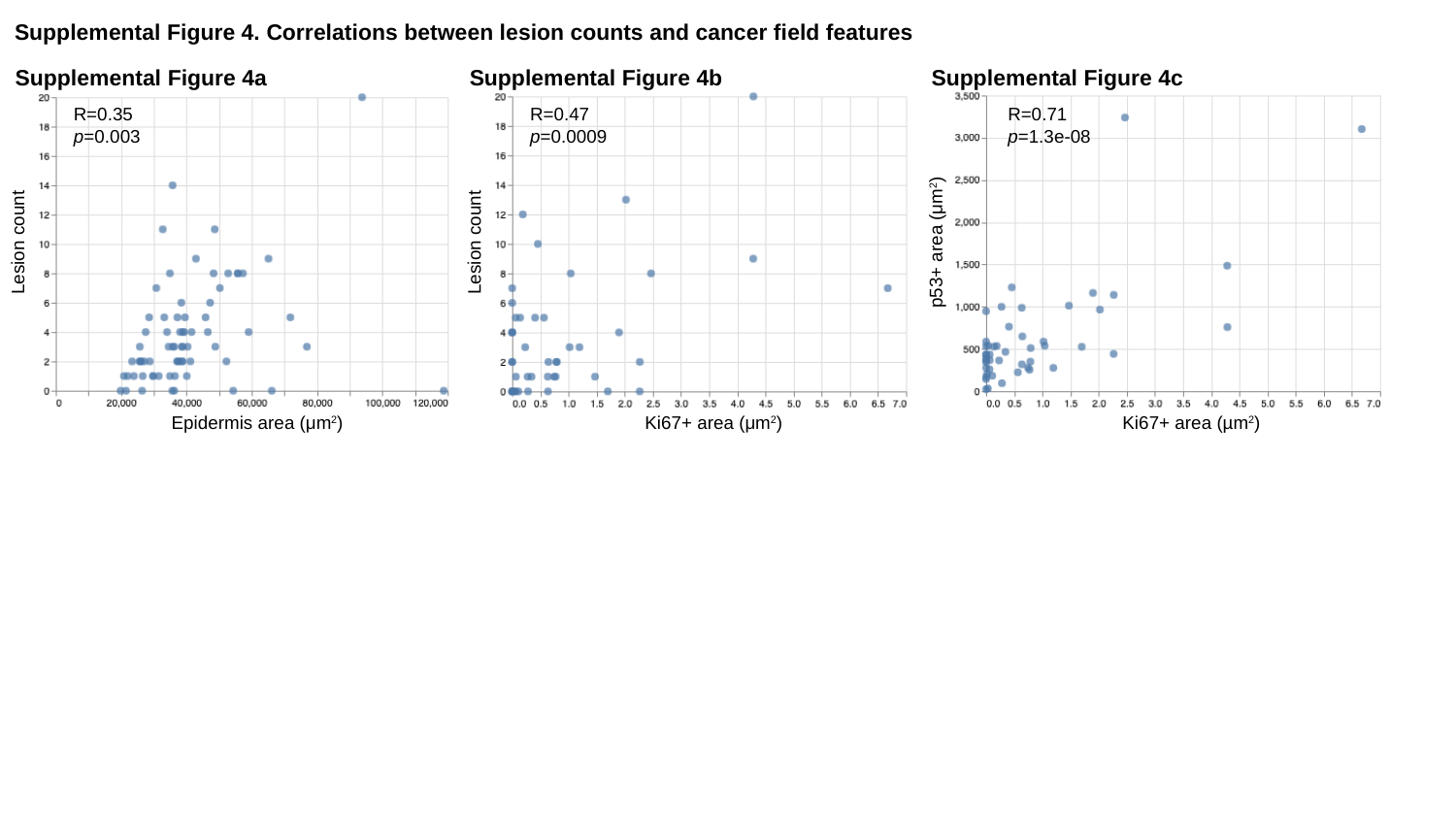

Supplemental Figure 4. Correlations between lesion counts and cancer field features
Supplemental Figure 4a
Supplemental Figure 4b
Supplemental Figure 4c
R=0.35
p=0.003
R=0.47
p=0.0009
R=0.71
p=1.3e-08
Lesion count
Lesion count
p53+ area (μm^2^)
Epidermis area (μm^2^)
Ki67+ area (μm^2^)
Ki67+ area (µm^2^)
